# *Salmonella* Enteritidis Effector AvrA suppresses autophagy by reducing Beclin-1 protein

**DOI:** 10.1101/2020.01.13.905596

**Authors:** Yang Jiao, Yong-guo Zhang, Zhijie Lin, Rong Lu, Yinglin Xia, Chuang Meng, Zhimin Pan, Xiulong Xu, Xinan Jiao, Jun Sun

## Abstract

Autophagy is a cellular process to clear pathogens. *Salmonella enterica* serovar Enteritidis (*S*.E) has emerged as one of the most important food-borne pathogens. However, major studies still focus on *Salmonella enterica* serovar Typhimurium. Here, we reported that AvrA, a *S*. Enteritidis effector, inhibited autophagy to promote bacterial survival in the host. We found that AvrA regulates the conversion of LC3 I into LC3 II and the enrichment of lysosomes. Beclin-1, a key molecular regulator of autophagy, was decreased after AvrA expressed strain colonization. In *S*.E-AvrA--infected cells, we found the increases of protein levels of p-JNK and p-c-Jun and the transcription level of AP-1. AvrA-reduction of Beclin-1 protein expression is through the JNK pathway. The JNK inhibitor abolished the AvrA-reduced Beclin-1 protein expression. Moreover, we identified that the AvrA mutation C186A abolished its regulation of Beclin-1 expression. In addition, AvrA protein interacted with Beclin-1. In organoids and infected mice, we explored the physiologically related effects and mechanism of AvrA in reducing Beclin-1 through the JNK pathway, thus attenuating autophagic responses.

**Importance:** *Salmonella* Enteritidis is an important pathogen with a public health concern and farm production risk, yet the host-pathogen interactions that govern the survival of *S*. Enteritidis infections are incompletely understood. Anti-bacterial autophagy provides potent cell-autonomous immunity against bacterial colonization. Here, we report that a new role for effector AvrA of *S*. Enteritidis in the reduction of Beclin-1 protein expression through the JNK pathway and the attenuation of the autophagic response in intestinal epithelial cells. This finding not only indicates an important role of *S*. Enteritidis effector in reducing host protein as a strategy to suppress autophagy, but also suggests manipulating autophagy as a new strategy to treat infectious diseases.

## Introduction

*Salmonella* is a Gram-negative, facultative anaerobe and an intracellular pathogen to both humans and animals. *Salmonella enterica* serovar Enteritidis has emerged as one of the most important food-borne pathogens for humans, and it is mainly associated with the consumption of contaminated poultry meat and egg (1, 2). Infection caused by *Salmonella* Enteritidis is the second most common cause of bacterial gastroenteritis in the developed world, and results in significant economic loss to the poultry industry and places a substantial burden on the healthcare system (2–4). *Salmonella* Enteritidis is an important pathogen with a public health concern, thus demanding further studying. However, the majority of basic researches in *Salmonella* field still focus on *Salmonella enterica* serovar Typhimurium (*Salmonella* Typhimurium) to study host-microbial interactions.

Autophagy pathway and proteins balance the beneficial and detrimental effects of immunity and inflammation, and thereby may protect against infectious (5). To survive in host cells, *Salmonella* use mechanisms to prevent clearance from host cells, such as escaping from the phagosome, inhibiting phagosome-lysosome fusion, and inhibiting apoptosis and autophagy in host cells (6–12). Among the numerous host defense systems against pathogens, anti-bacterial autophagy provides potent cell-autonomous immunity against bacterial attempts to colonize the cytosol of host cells (5, 13–15). During this process, the phagophore expands and engulfs pathogens, and closes to originate the autophagosome that fuses with the lysosome, at which the degradation of the pathogens takes place (16). There are more than 20 ATG proteins (many of which are evolutionarily conserved) that are essential for the execution of autophagy (17). Notably, the mammalian autophagy protein Beclin-1, an ortholog of the Atg6 in yeast, is a key molecule regulator of autophagy. Beclin-1 interacts with several cofactors (e.g. Atg14L, HMGB1, IP3R, PINK, and survivin) to regulate the lipid kinase Vps-34 protein and promote the formation of Beclin-1-Vps34-Vps15 core complexes, thereby inducing autophagy (18, 19).

*Salmonella* possesses a range of effector proteins that are translocated into the host cells via a type III secretion system (T3SS). These effector proteins are generally assumed to influence the host’s cellular functions to facilitate *Salmonella* invasion and intracellular carriage (20–23). AvrA is one of the *Salmonella* effectors secreted by the *Salmonella* pathogenicity island 1 (SPI-1) T3SS. The AvrA protein in *Salmonella* Typhimurium is an anti-inflammatory effector that possesses acetyltransferase activity and inhibits the host c-Jun N-terminal kinase (JNK)/AP-1 and NF-κB signaling pathways. Through these methods, AvrA inhibits the host inflammatory response and stabilizes the intestinal tight junctions to the benefit of bacterial survival (9, 24–28). However, the role of AvrA in *Salmonella* infection and host autophagic response is unexplored.

Here, we hypothesize that *Salmonella* Enteritidis effector AvrA inhibits the autophagic response by decreasing Beclin-1 expression. We used wild-type, *Salmonella* Enteritidis C50336 and established a deletion *Salmonella* Enteritidis mutant *S*.E-AvrA^-^ and a plasmid-mediated complementary strain *S*.E-AvrA^-^/pAvrA^+^(*S*.E-AvrA^+^) (28). In cell cultures, organoids and infected mice, we explored the physiologically related effects and molecular mechanism of AvrA regulation of autophagy in intestinal epithelial cells, whereas most studies on bacterial effectors and autophagy only use cell cultures. Our study provides new insights into the mechanisms of the bacterial effects in regulating host-microbial interactions.

## Results

### *Salmonella* Enteritidis AvrA decreases autophagy markers and enhanced bacterial invasion in vitro

Autophagy is an important cell-autonomous defense mechanism required for pathogen clearance (13, 14). LC3 and p62 are well-recognized markers for autophagic activity (29). In this study, we found that in human intestinal epithelial HCT116 cells, the conversion of LC3 I into LC3 II was increased after *S*.E-AvrA^-^ infection compared to that after wild-type *S*.E or *S*.E-AvrA^+^ infection. Notably, p62, which is a bona fide target of autophagosomal degradation, was decreased in the cells infected with the *S*.E-AvrA mutant strain compared with the expression in cells infected with the *S*.E-WT or *S*.E-AvrA^+^ strains (Fig. 1A). A densitometry analysis showed a significant difference between the cells infected with the different *S*.E strains (Fig. 1B, C). Meanwhile, HCT116 cells pre-treated with LysoTracker showed more lysosomes in the cells infected with the *S*.E-AvrA mutant strain than in the cells infected with the AvrA expressed strains (Fig. 1D, E). The role of AvrA in *Salmonella* Enteritidis invasion is unknown. We further compared the invasion ability of *Salmonella* Enteritidis strains with or without AvrA expression. We found that *S*.E-AvrA^-^ colonized human epithelial cells showed a decreased intracellular bacterial load compared to those colonized with wild type *S*.E or *S*.E-AvrA^+^ (Fig. 1F). We observed similar trends in autophagic activity following *S*.E-AvrA^-^, *S*.E-AvrA^+^ and *S*.E-WT infection in human Caco-2 BBE and SKCO-15 cells (data not shown). Taken together, these data suggest that the *Salmonella* Enteritidis effector AvrA inhibits autophagy i*n vitro*.

**Figure 1.**
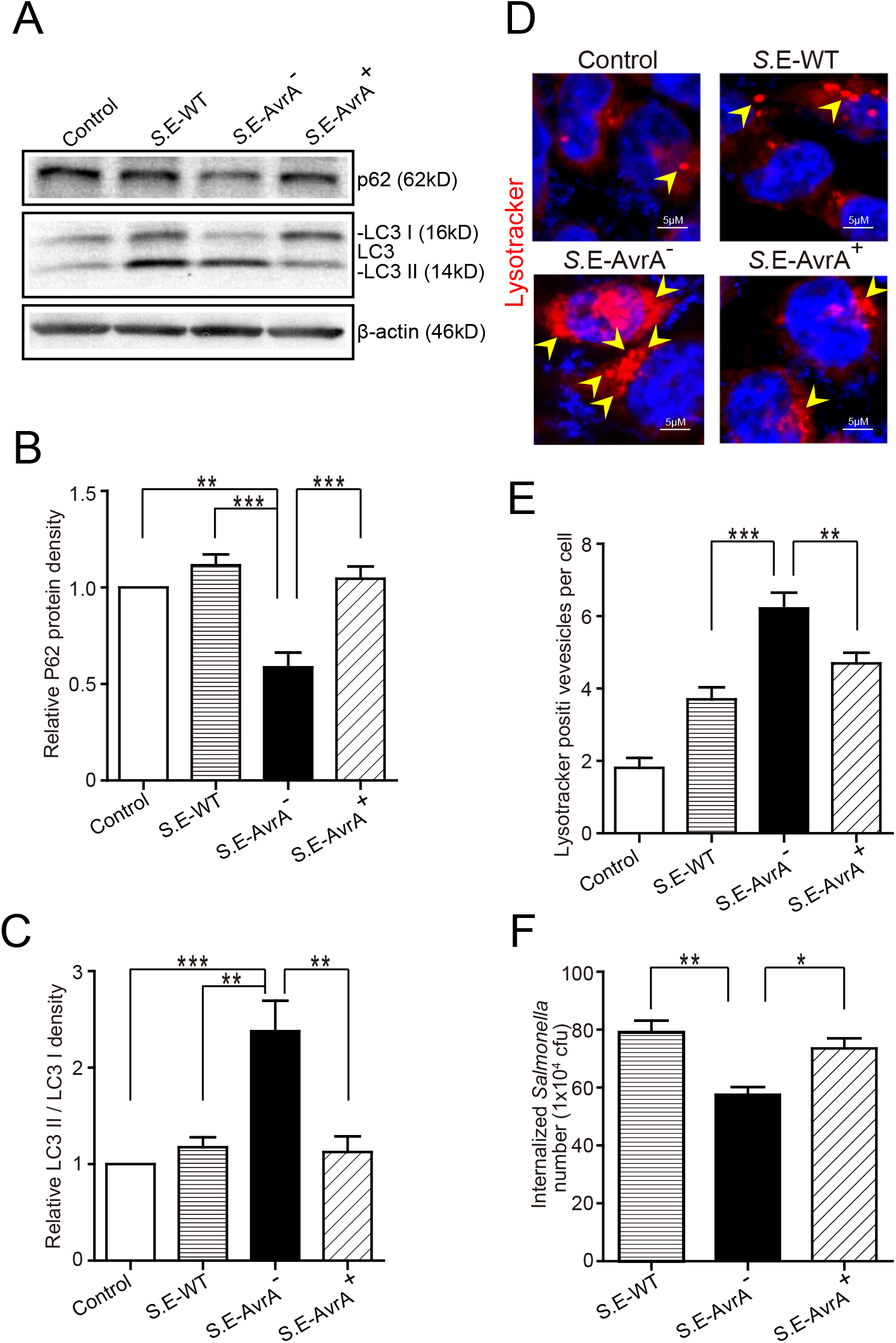
*Salmonella* Enteritidis AvrA inhibits autophagy in cell models. AvrA-regulated expression levels of autophagy related proteins. **(A)** The indicated HCT116 cell lines were infected with *S*.E WT, *S*.E-AvrA^-^ and *S*.E-AvrA^+^ strains (MOI:1:1, 1h incubation before harvested, n=4) as shown and analyzed for protein expression by immunoblotting. The immunoblotting of P62/LC3 was used to track the expression of P62 and the conversion of LC3 I into LC3 II for autophagic activity in the HCT116 cells after infection with the different *S*.E. The relative density of P62 **(B)** and LC3 II/LC3 I **(C)** was determined using Quantity One 4.6.2 software (Bio-Rad, Hercules, CA, USA). n=4, **adjusted *P* < 0.01, ***adjusted *P* < 0.001 by ANOVA test. **(D)** HCT116 cell lines were incubated with 100 nM LysoTracker Deep Red Probe, then infected with S.E WT, S.E-AvrA- and S.E-AvrA+ strains (MOI:1:1, 1h incubation before microscopic examination) to check the lysosomes staining. The immunofluorescence indicated that the HCT116 cells, pre-treated with LysoTracker, showed more lysosomes in the cells infected with the *S*.E-AvrA^-^ bacteria compared with the cells infected by the AvrA present strains. **(E)** Quantification of the number of lysotracker positive vesicles. The data are reported as the mean ± SE from three independent experiments, and a total of 100 cells per condition were analyzed. n=100, **adjusted *P* < 0.01, ***adjusted *P* < 0.001 by ANOVA test. **(F)** The numbers of internalized *Salmonella* in human epithelial cells colonized with wild-type *Salmonella* Enteritidis or AvrA mutant or AvrA-complemented strains. Human epithelial cells were grown on an insert, colonized with an equal number of the indicated bacteria for 30 min, after washed with HBSS, cells were incubated in DMEM containing gentamicin (100 μg/ml) for 30 min, and the number of internalized *Salmonella* (*Salmonella* invasion) was then determined. Data are reported as the mean ± SE from six independent experiments, *adjusted *P* < 0.05, **adjusted *P* < 0.05 by ANOVA test.

### AvrA reduces Beclin-1 at the protein levels and interacts with Beclin-1

Beclin-1, a key molecular regulator of autophagy, interacts with several cofactors to regulate the lipid kinase Vps-34 protein and promote the formation of Beclin-1-Vps34-Vps15 core complexes, thereby inducing autophagy (18, 19). Thus, we determined whether the protein level of Beclin-1 was changed by the infection with the different *S*.E strains. As shown in Fig. 2A and 2B, Beclin-1 protein expression was significantly decreased after colonization of the AvrA present strains for 1 hour. In contrast, the cells colonized with the *S*.E-AvrA^-^ bacteria maintained Beclin-1 protein expression. To verify that AvrA affects the protein expression of Beclin-1, we transfected an AvrA WT plasmid or an AvrA C186A mutant [mutated at the key cysteine required for AvrA activity (24)] plasmid into HCT116 cells. As expected, the AvrA WT plasmid decreased endogenous Beclin-1 protein expression. However, the AvrA C186A mutant plasmid maintained the endogenous Beclin-1 protein expression (Fig. 2C, D,). We further determined the interaction of AvrA/Beclin-1 in the HCT116 cells by immunoprecipitation. Vps34 was used as a positive control. We found that exogenous AvrA (c-myc tag) coimmunoprecipitated with exogenous Beclin-1 (HA-tag), suggesting that AvrA interacted with Beclin-1 (Fig. 2E). Therefore, the data suggest that the *S*.E effector AvrA interacts with Beclin-1 and changes the Beclin-1 protein level to inhibit autophagy.

**Figure 2.**
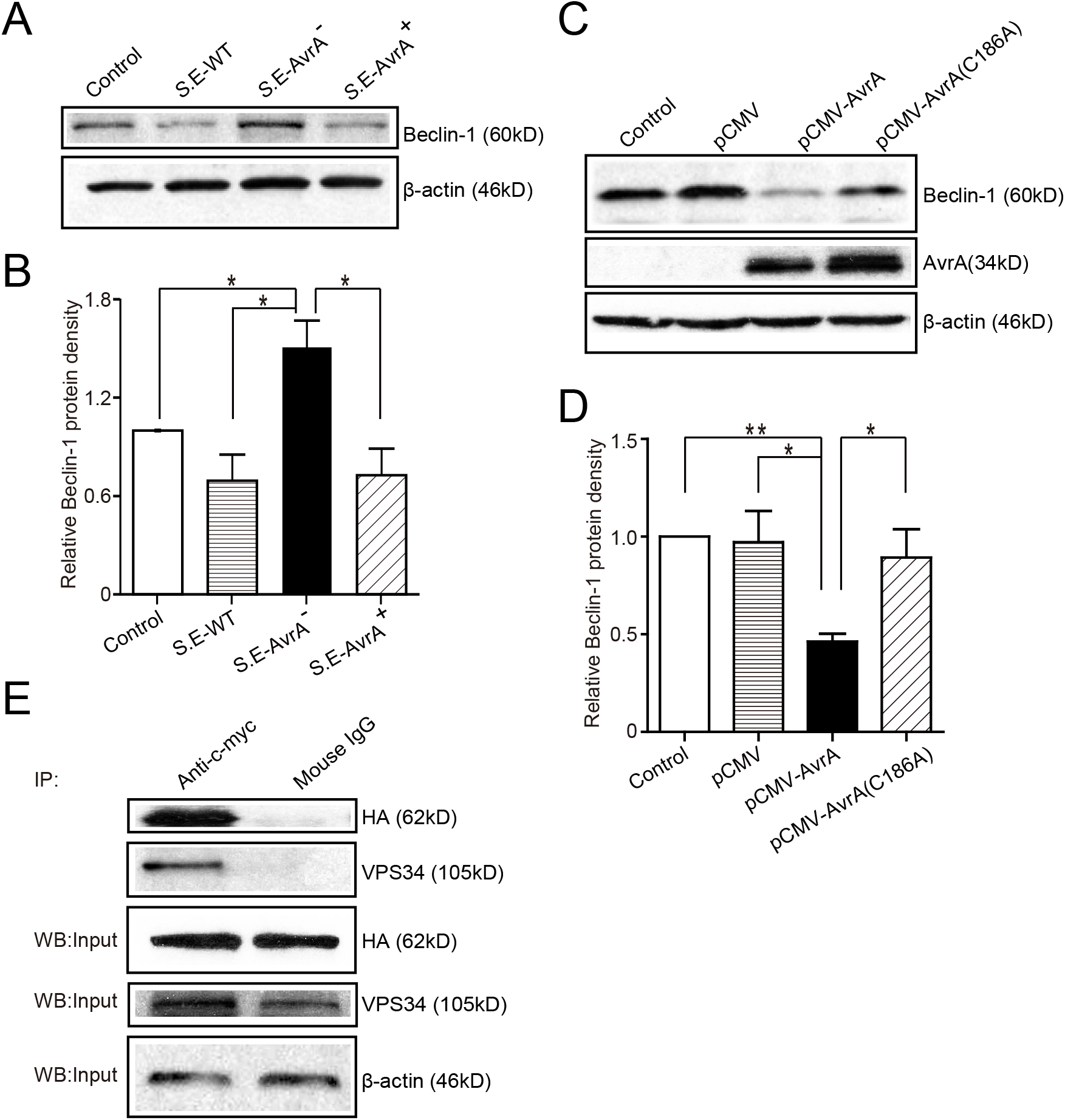
AvrA changes Beclin-1 protein levels and interacts with Beclin-1 in HCT116 cells. AvrA changes Beclin-1 protein levels and interacts with Beclin-1 in HCT116 cells. **(A)** The Western blot shows the expression of Beclin-1 in the HCT116 cells after colonization with wild-type *Salmonella* Enteritidis or AvrA mutant or AvrA-complemented strains (MOI:1:1, 1h incubation before harvested, n=4). The relative density of Beclin-1 **(B)** was determined using Quantity One 4.6.2 software (Bio-Rad, Hercules, CA, USA). n=4, *adjusted *P* < 0.05 by ANOVA test. **(C)** The Western blot shows the expression of Beclin-1 in the HCT116 cells after transfection with the AvrA wild-type and AvrA mutant plasmids (200 ng/μl, 24h incubation, n=5). The relative density of Beclin-1 **d** was determined using Quantity One 4.6.2 software (Bio-Rad, Hercules, CA, USA). n=5, *adjusted *P* < 0.05, **adjusted *P* < 0.01 by ANOVA test. **(E)** The HCT116 cells were cotransfected with the AvrA WT plasmid (c-myc tag) and the Beclin-1 WT plasmid (HA-tag) (200 ng/μl, 24h incubation, n=4). At the indicated times, immunoprecipitation was performed with an anti-c-myc mouse monoclonal antibody. Pre-immune mouse IgG was used as a negative control. VPS34 was used as a positive control. The Western blot analyses of the pre-immunoprecipitation (Input) and immunoprecipitated samples (IP) were performed with an anti-c-myc mouse monoclonal antibody or with an anti-HA mouse monoclonal antibody. These results shown are representative of three independent experiments.

### AvrA inhibits the JNK signaling pathway to decrease Beclin-1

Beclin-1 is regulated by the JNK signaling pathway (30). Previous studies have shown that *Salmonella* AvrA inhibits the activation of the JNK signaling pathways (9, 28). Using Western blotting, we found that the protein levels of p-JNK and p-c-Jun were higher in the *S*.E-AvrA^-^-infected cells than in the cells infected by the *S*.E-WT or *S*.E-AvrA^+^ strains (Fig. 3A). Meanwhile, a luciferase reporter assay showed that AP-1 transcription was increased, as a consequence of the activation of JNK/c-JUN (Fig. 3B). These data suggest that the JNK/c-jun pathway and AP-1 transcription are more highly activated in the *S*.E-AvrA^-^-infected cells than in the cells infected with the AvrA present strains. Interestingly, after treatment with the JNK inhibitor SP600125, the level of Beclin-1 and P62 protein expression was not different between the cells infected with AvrA or without AvrA (Fig. 3A, B), indicating that the AvrA-related responses were abolished by the JNK inhibitor. Thus, our data indicate that the *S*.E effector AvrA inhibits the autophagic response by decreasing Beclin-1 at the protein level, and it occurs by inhibiting the JNK/c-Jun/AP-1 signaling pathways.

**Figure 3.**
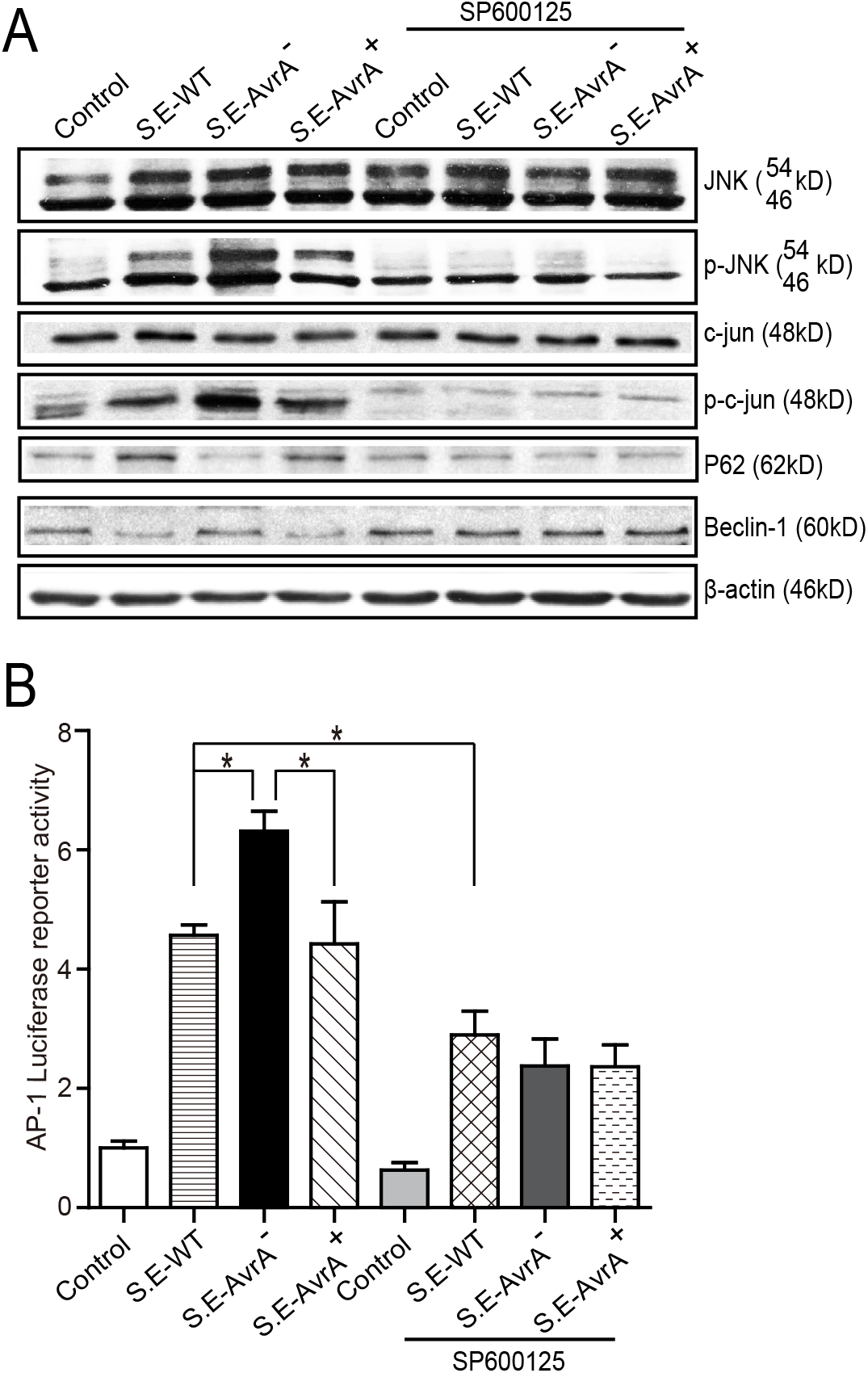
AvrA inhibits the JNK/c-jun/AP-1 signaling pathway to decrease Beclin-1 in HCT116 cells. AvrA inhibited the JNK pathway and decreased Beclin-1 in HCT116 cells, and the effect of AvrA on Beclin-1 expression was abolished by the JNK inhibitor SP600125. **(A)** The Western blot shows a change in the JNK pathway markers and Beclin-1 in the cells treated with the JNK inhibitor SP600125 (50 μM, 30 min) and those infected with the different *S*.E strains (MOI:1:1, 1h incubation before harvested, n=4), These results shown are representative of four independent experiments. **(B)** The luciferase reporter assay shows a change in AP-1 transcription in the cells treated with the JNK inhibitor SP600125 and in those infected with the different *S*.E strains (MOI:1:1, 1h incubation before harvested, n=6). *adjusted *P* < 0.05 by ANOVA test.

### The AvrA mutant C186A plasmid expression abolishes the regulation of exogenous Beclin-1 expression

To further study the function of the AvrA protein, we cotransfected an AvrA WT plasmid or an AvrA C186A mutant plasmid with a Beclin-1 WT plasmid. The AvrA C186A mutation is known to abolish the enzyme activity of AvrA (24). Our data showed that the AvrA WT plasmid decreased not only the endogenous Beclin-1 protein but also the exogenous Beclin-1 protein. In contrast, the AvrA C186A mutant plasmid abolished its regulation on Beclin-1 expression (Fig. 4A, B). Moreover, we found that the protein levels of p-JNK, p-c-Jun and Beclin-1 were decreased in the AvrA WT plasmid transfected cells compared with the cells transfected with the AvrA C186A mutant plasmid. These data verified that AvrA decreased Beclin-1 by inhibiting the JNK signaling pathways, whereas the AvrA C186A mutation abolished its regulation (Fig. 4C). We did not observe the change of the Beclin-1 at the mRNA level (data not shown).

**Figure 4.**
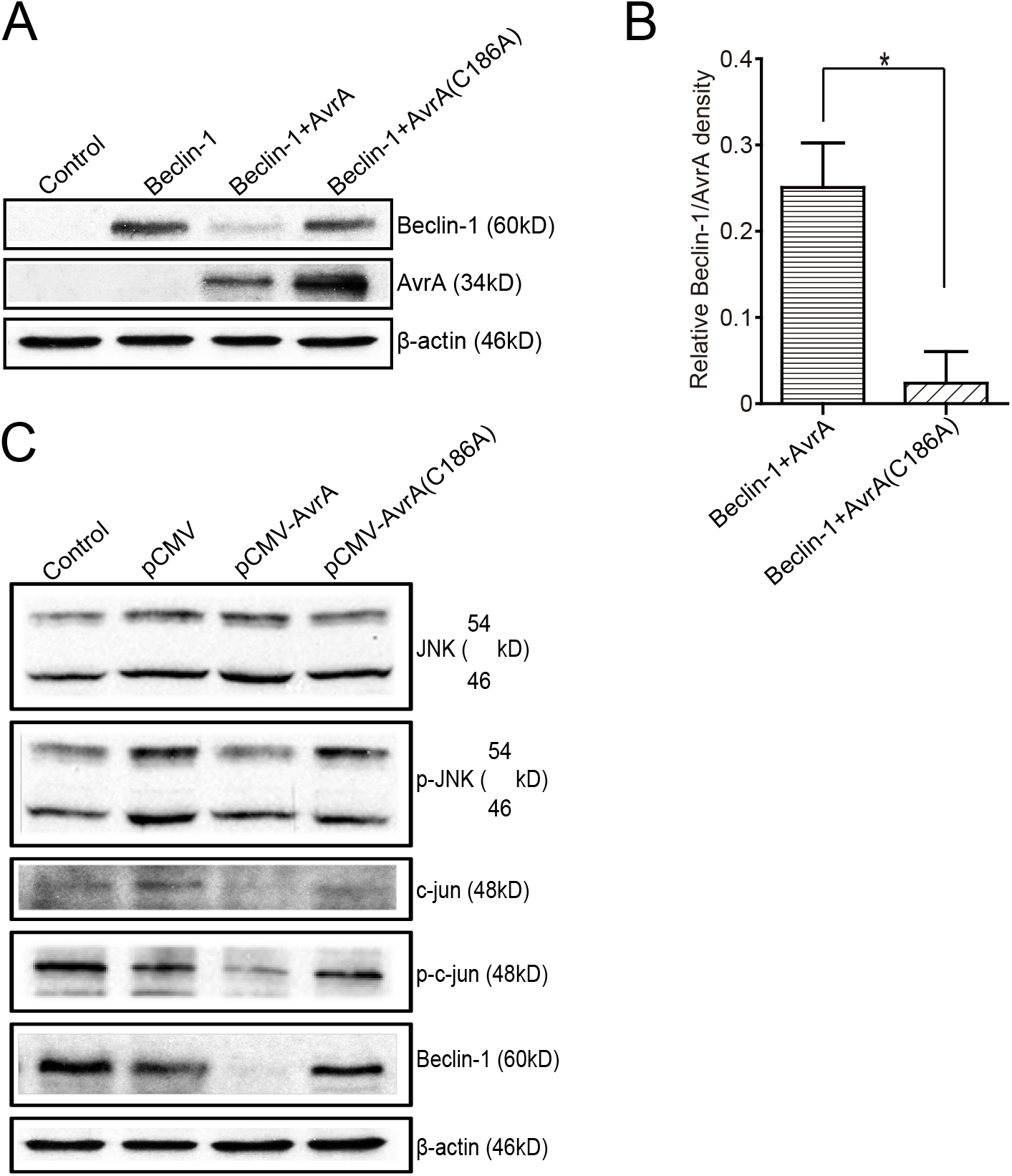
The AvrA C186A mutant plasmid abolishes the regulation of exogenous Beclin-1. The HCT116 cells were transfected with the indicated plasmids (200 ng/μl, 24h incubation, n=4). At the indicated times, the changes in the target proteins were measured. **(A)** The Western blot shows the expression of Beclin-1 in the HCT116 cells after transfection with the indicated plasmids. The relative density of Beclin-1 **(B)** was determined using Quantity One 4.6.2 software (Bio-Rad, Hercules, CA, USA). n=4, \**P* <0.05 by student’s t-test. **(C)** The Western blot shows the change in the JNK pathway markers and Beclin-1 in the HCT116 cells after transfection with the AvrA wild-type and AvrA mutant plasmids. These results shown are representative of four independent experiments.

### AvrA expressing bacteria reduce the level of Beclin-1 protein in mouse organoids

Intestinal organoid culture is a newly developed 3D system to determine the bacterial–epithelial interactions post *Salmonella* infection (31). We found that the Beclin-1 protein expression was significantly decreased after the infection of the *S*.E-WT expressing AvrA. In contrast, the organoids colonized with the *S*.E-AvrA^-^ mutant strain increased Beclin-1 protein expression, whereas *S*.E-AvrA^+^ could reduce the Beclin-1 expression (Fig. 5A, B). Moreover, we found AvrA-associated changes of P62, LC3 II / LC3 I, and p-JNK in the *Salmonella*-infected organoids (Fig. 5A, C, D). Decreased autophagy marker P62 was further confirmed in organoids colonized with the *S*.E-AvrA^-^ strain in comparison with organoids infected by the *S*.E-WT and *S*.E-AvrA^+^ strains by Immunostaining (Fig. 5E). These changes in the 3D organoids were consistent with AvrA-suppressed autophagy observed in the 2D cultured cell lines.

**Figure 5.**
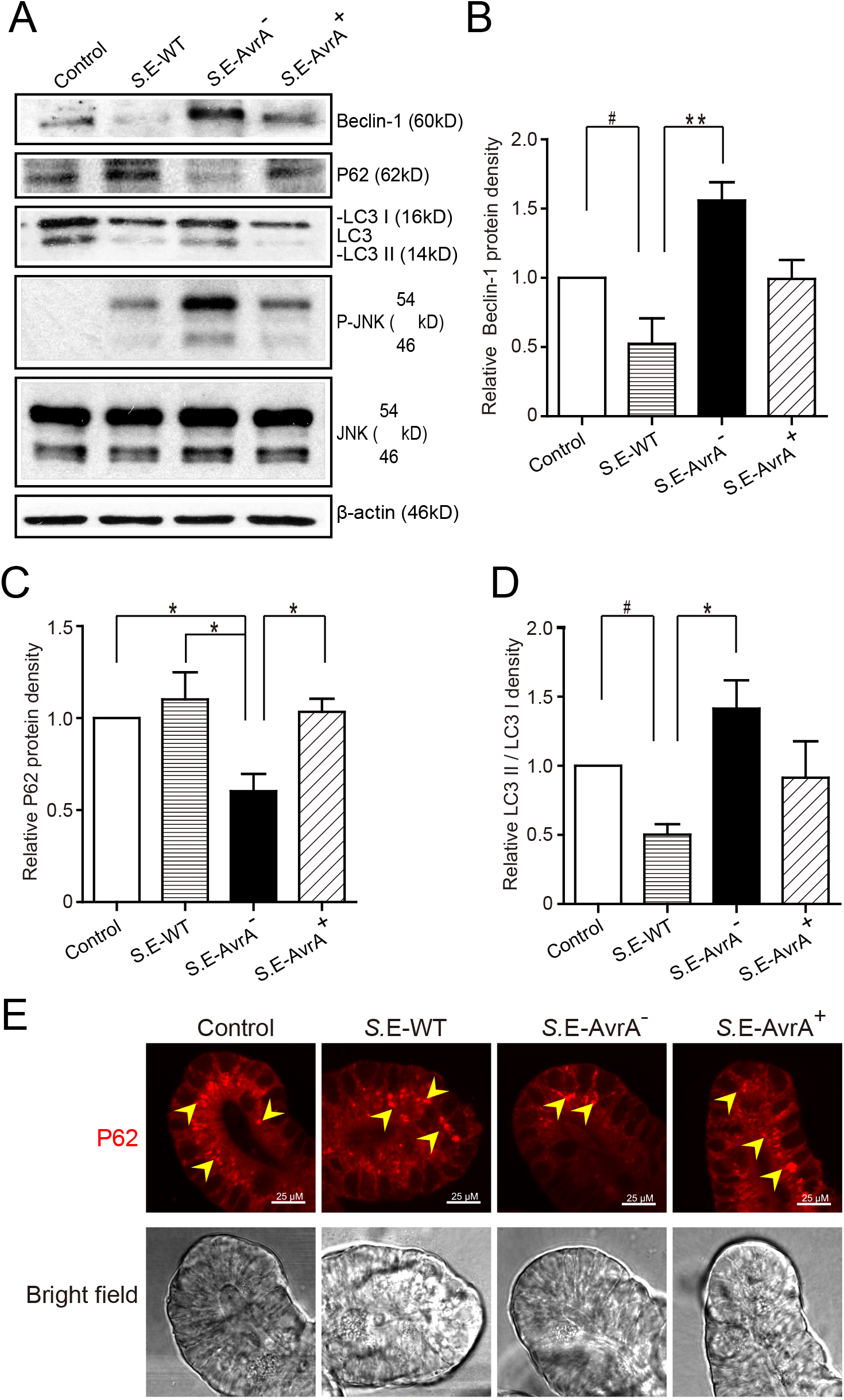
*Salmonella* Enteritidis AvrA changes the levels of Beclin-1 in mouse intestinal organoids. *Salmonella* Enteritidis infection and invasion in the mouse organoids. **(A)** The Western blot shows the expression of Beclin-1, P62, LC3 and the JNK pathway markers in the organoids after infection with wild-type *Salmonella* Enteritidis or AvrA mutant or AvrA-complemented strains (MOI:1:1, 1.5h incubation before harvested, n=3). The relative density of Beclin-1 **(B)**, P62 **(C)** and LC3 II/LC3 I **(D)** was determined using Quantity One 4.6.2 software (Bio-Rad, Hercules, CA, USA). n=3, *adjusted *P* < 0.05, **adjusted *P* < 0.01, #unadjusted *P* < 0.05 by ANOVA test. **(E)** The representative images of the immunostaining of P62 in the organoids after infection with wild-type *Salmonella* Enteritidis or AvrA mutant or AvrA-complemented strains (MOI:1:1, 1.5h incubation).

### AvrA changes the levels of Beclin-1 and affects the function of Paneth cell granules of the ileal tissues in a mouse model

To study the role of the *S*.E effector AvrA in an *in vivo* model of natural intestinal infection, we used the streptomycin pretreatment mouse model of enteric salmonellosis (32, 33). C57BL/6 mice (female, 6-8 weeks) were pretreated with streptomycin for 24 hours before infection with the *S*.E-WT, *S*.E-AvrA^-^ or *S*.E-AvrA^+^ strains by oral gavage. In the Ileum samples from the *S*.E-WT-infected mice, Beclin-1 protein expression was significantly decreased compared to the expression in the samples from the *S*.E-AvrA^-^-infected mice (Fig. 6A, B). As expected, decreased P62, increased conversion of LC3 I into LC3 II and activation of the JNK pathway were also found in mice infected with the *S*.E-AvrA^-^ strain compared with those in the mice infected by the *S*.E-WT strain (Fig. 6A, C, D). These data suggest that AvrA inhibits the JNK signaling pathway to decrease Beclin-1 expression and impair the autophagic response *in vivo*.

**Figure 6.**
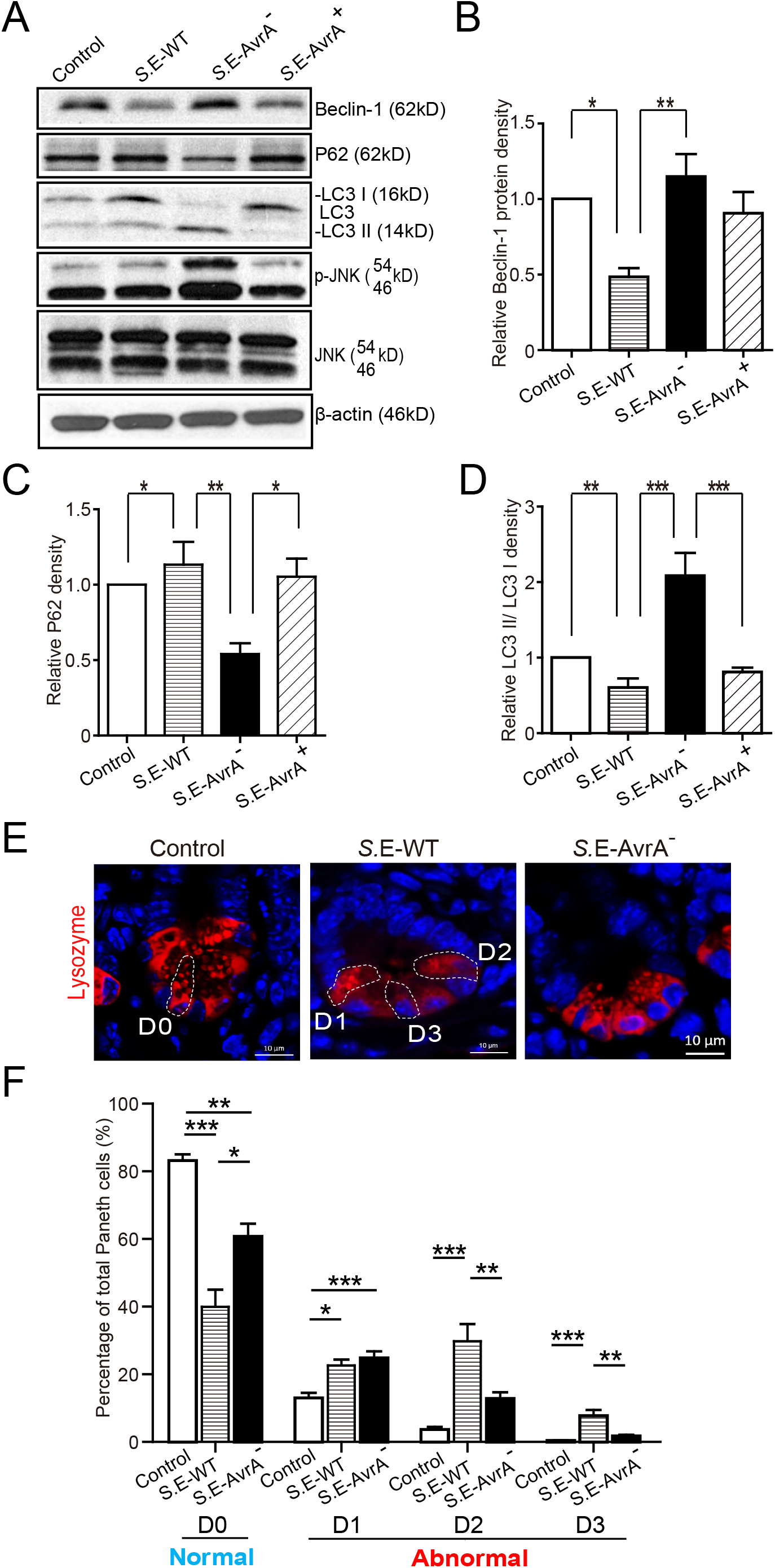
AvrA changes the levels of Beclin-1 and affects the function of Paneth cell granules of the ileal tissues in a mouse model. Wild-type *Salmonella* Enteritidis, AvrA mutant or AvrA-complemented strains were used to infect C57BL/6 mouse models (1.0×10^8^ CFU per mouse, i.g., n=6). At 8h postinfection, the ileal tissue was harvested for immunoblotting and immunofluorescence staining. **(A)** Western blot shows the expression of Beclin-1, P62, LC3 and the JNK pathway markers in the ileal tissue after infection with the different *S*.E strains. The relative density of Beclin-1 **(B)**, P62 **(C)** and LC3 II/LC3 I **(D)** was determined using Quantity One 4.6.2 software (Bio-Rad, Hercules, CA, USA). n=5-6, *adjusted *P* < 0.05, **adjusted *P* < 0.01 by ANOVA test. **(E)** The representative images of the indirect immunofluorescence of the sections stained for lysozymes (red) in the ileal crypts of the C57BL/6 mice following infected with wild-type *Salmonella* Enteritidis or AvrA mutant strains. **(F)** The percentage of Paneth cells displaying a normal (D0) and abnormal (D1 to D3) pattern of lysozyme expression. n=5-6, *adjusted *P* < 0.05, **adjusted *P* < 0.01, ***adjusted *P* < 0.001 by ANOVA test.

Deficits in the autophagy pathway impair Paneth cell function in intestine (34, 35). Thus, we counted the number of Paneth cells using a previously reported method to stain lysozymes (35, 36). The abnormal Paneth cells were grouped as D1 (disordered), D2 (depleted), or D3 (diffuse) (Fig. 6E). We found fewer normal Paneth cells (D0) in the mice infected with the *S*.E-WT strain than in the mice infected with the *S*.E-AvrA^-^ strain (Fig. 6E, F). Consequently, the number of abnormal Paneth cells (D1-D3) increased in the mice infected with the wild-type *S*.E strain (Fig. 6F).

## Discussion

In the current study, we report that the *S*. Enteritidis effector protein AvrA decreased Beclin-1 protein expression, thus impairing the autophagic response, for the benefit of *Salmonella* survival. AvrA-mediated regulation of host autophagy involves blocking the JNK signaling pathway, which was demonstrated *in vitro* and *in vivo* (Fig. 7E). The *S*.Enteritidis effector protein AvrA inhibited the JNK signaling pathways in epithelial cells, impaired autophagy by decreasing Beclin-1 expression at the protein level *in vitro* and *in vivo*, and affected the function of Paneth cell granules in a mouse model. Moreover, the JNK inhibitor SP600125 abolished the AvrA-reduced Beclin-1 protein expression. Together, these data suggest that the *S*.E effector AvrA inhibits the JNK/c-Jun/AP-1 signaling pathway to decrease Beclin-1 expression. In this way, *Salmonella* Enteritidis impairs the autophagic response to the benefit of the pathogen’s survival. Moreover, AvrA affected the function of Paneth cell granules, potentially by inhibiting autophagy (Fig. 7E).

**Figure 7.**
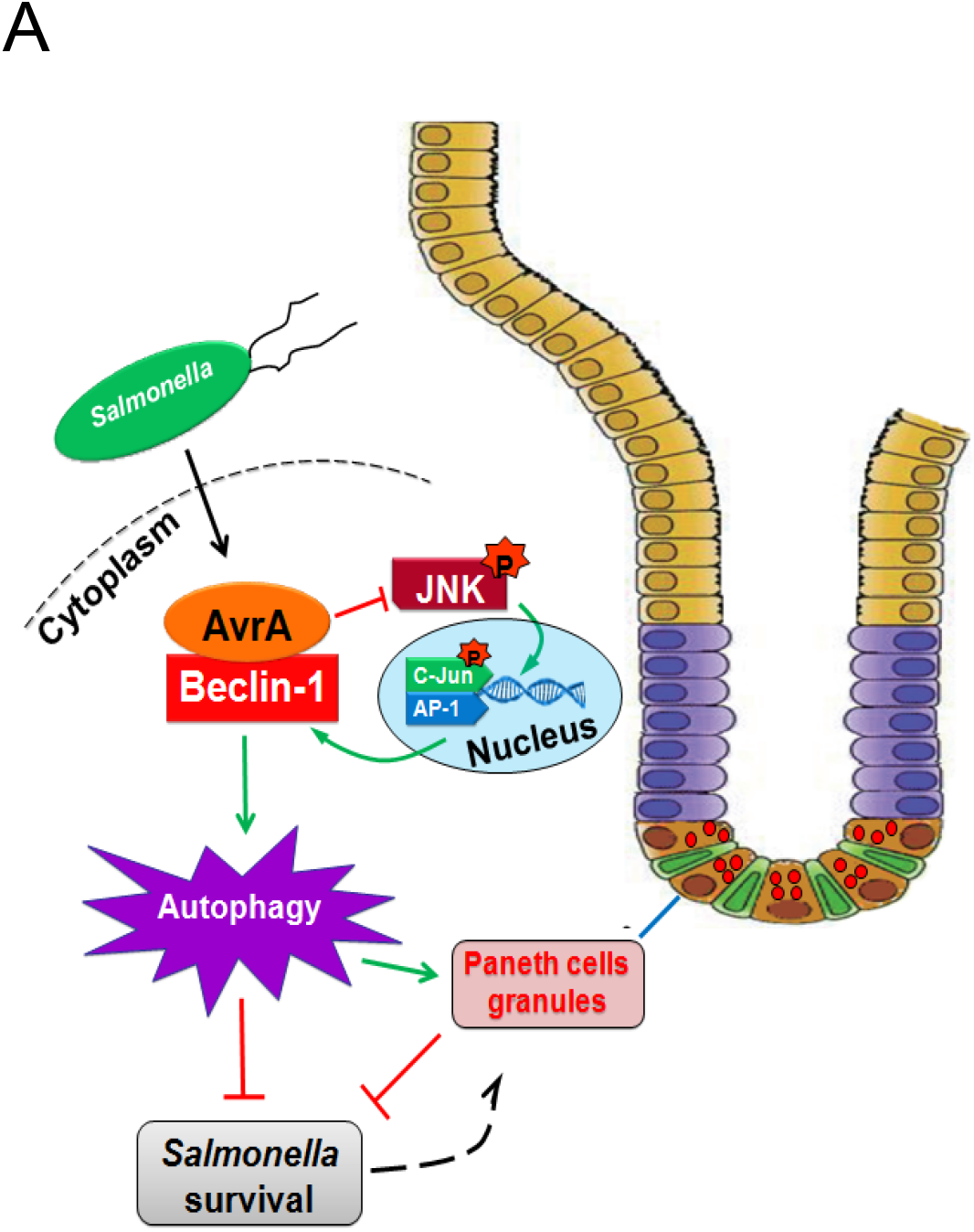
Working model of AvrA inhibition of Beclin-1-dependent autophagy. *Salmonella* Enteritidis AvrA inhibits the autophagic response by decreasing Beclin-1 protein. At the molecular level, AvrA interacts with Beclin-1. This process occurs through the inhibition of the JNK signaling pathway. At the cellular level, AvrA affects the function of Paneth cell granules by inhibiting autophagy.

The AvrA from *S*. Typhimurium is known as an anti-inflammatory effector that possesses acetyltransferase activity toward specific host MAPKKs and inhibits the host JNK/c-Jun/AP-1 and NF-κB pathways, but the role of AvrA from *S*. Enteritidis is less explored (9, 24-28). Here, we demonstrated that *S*. Enteritidis AvrA inhibited the autophagic response. Beclin-1 protein levels were reduced with AvrA present strain infection. In contrast, *S*.E-AvrA mutant strain infection maintained Beclin-1 protein expression. Earlier studies on the mechanism underlying the regulation of autophagy in cancer cells showed that the activation of the JNK pathway may be involved in the regulation of Beclin-1 expression, and the latter event could be responsible for the induction of the autophagic response (30). Blocking the JNK pathway might be the reason that AvrA decreases Beclin-1 expression. To verify our results, we used a specific JNK inhibitor SP600125 to block the JNK signaling pathway. After treatment with the JNK inhibitor SP600125, Beclin-1 protein expression did not differ between infection with and without AvrA. The results demonstrated that the *S*. Enteritidis effector AvrA inhibits the JNK/c-Jun/AP-1 signaling pathway to decrease Beclin-1 expression. Our data indicated that cysteine 186 is the key amino acid required for the AvrA regulation of Beclin-1 expression. We further demonstrated that the single mutation of Cys^186^ blocked JNK activity and abolished the AvrA-induced downregulation of Beclin-1. Thus, we have demonstrated a strategy that *Salmonella* Enteritidis used to impair the autophagic response to the benefit of the pathogen’s survival.

In the current study, we observed that the *S*.E WT strain with AvrA expression and complementary strain induced weak autophagic activity, compared to the autophagic activity following *S*.E AvrA muant strain infection. Numerous lines of evidence indicate that *Salmonella* infection can activate robust host cell autophagy (37–39). This difference maybe due to the serotype difference. The existing research tends to use *Salmonella* Typhimurium to study the interaction between host autophagic activity and bacterial invasion. However, *Salmonella* Enteritidis, which belongs to another *Salmonella* serotype, could be much difference from *Salmonella* Typhimurium in host-adaptability, virulence, intracellular survival and so on (40–42).

Interestingly, our data showed that AvrA interacted with Beclin-1 and decreased the ubiquitination of Beclin-1 (Fig. S1). However, the detailed mechanism is unclear. Previous studies have shown that the ubiquitination of Beclin-1 enhances its association with Vps34 to promote Vps34 activity (43). Thus, AvrA may suppress Beclin-1 ubiquitination to inactivate Vps34 activity, leading to the suppression of autophagy. Certainly, this hypothesis requires further study.

Here, we highlighted the organoid system for investigating the host-bacterial interactions. Our previous studies have demonstrated that the intestinal organoid culture is a newly developed 3D system to study *Salmonella* infection (31, 44). Our data in the organoid system have shown that AvrA expressing bacteria reduce the levels of Beclin-1, thus suppressing autophagy in the host. In the future, human organoids could be used to further understand how bacterial effectors manipulate the host responses.

Paneth cells are specialized epithelial cells that are primarily located in the small intestine. The granules of Paneth cells contain AMPs-α-defensins, lysozyme, and secretory phospholipase A2 (45–47). Among these, lysozyme is a useful marker of the Paneth cell secretory granule (48). Our data showed that the normal expression pattern of the Paneth cells decreased in the mice infected with the AvrA present strain compared with that in the mice infected with the *S*.E-AvrA mutant strain. The ileum, after infection with the AvrA present strain, contained an increased proportion of Paneth cells with disorganized or diminished granules or exhibited diffuse cytoplasmic lysozyme staining. These findings suggest that AvrA impairs the autophagic response, and then, the autophagy affects the biosynthesis or quality control of the lysosomal pathway in the Paneth cell granules. Thus, there were fewer normal Paneth cells in the mice infected with the *S*.E-WT strain. Recently, a study showed that *S*. Typhimurium invaded Paneth cells, and the invasion was associated with an elevated number of LC3^+^ puncta in the Paneth cells (49). It is not clear whether *S*. Enteritidis interact with Paneth cells and affect autophagy through the AvrA effector. Alternatively, the Paneth cells with fewer granules might be a result of Paneth cell exhaustion (34), compensating for the changes elsewhere in the epithelium, due to the survival benefit of S*almonella* caused by AvrA suppressing autophagy.

Taken together, our data reveal a new role for AvrA in *S*. Enteritidis in the reduction of Beclin-1 protein expression through the JNK pathway and the attenuation of the autophagic response in intestinal epithelial cells. Bacterial effector proteins paralyze or reprogram host cells to the benefit of the pathogens. Our findings indicate an important role of *S*. Enteritidis effector in reducing host protein as a strategy to suppress autophagy. Manipulating autophagic activity through the JNK pathway may be a novel therapeutic approach to treat infectious diseases.

## Materials and Methods

### Animals and Ethics Statement

C57BL/6 mice (female, 6-8 weeks) were obtained from the Jackson Laboratory (Jackson Laboratory, Bar Harbor, ME, USA). All the animal work was approved by the University of Illinois at Chicago Committee on Animal Resources. Euthanasia was accomplished via sodium pentobarbital administration (100 mg per kg body weight, i.p.) followed by cervical dislocation. All procedures were conducted in accordance with the approved guidelines of the Committees on Animal Resources.

### Bacterial strains and growth conditions

The wild-type *Salmonella* Enteritidis strain C50336 (*S*.E WT) was obtained from the National Institute for the Control of Pharmaceutical and Biological Products (NICPBP), China. The *Salmonella* Enteritidis AvrA mutant strain *S*.E-AvrA^-^ was derived from C50336, and the AvrA complemented strain *S*.E-AvrA^+^ was constructed as in previous studies by our laboratory (28). The bacteria and plasmids used in this study are listed in **Table I**. The bacterial growth conditions were as follows: the non-agitated microaerophilic bacterial cultures were prepared by inoculating 10 ml of Luria-Bertani (LB) broth with 0.01 ml of a stationary phase culture followed by an overnight incubation (>18 hours) at 37°C (50).

**Table I.**
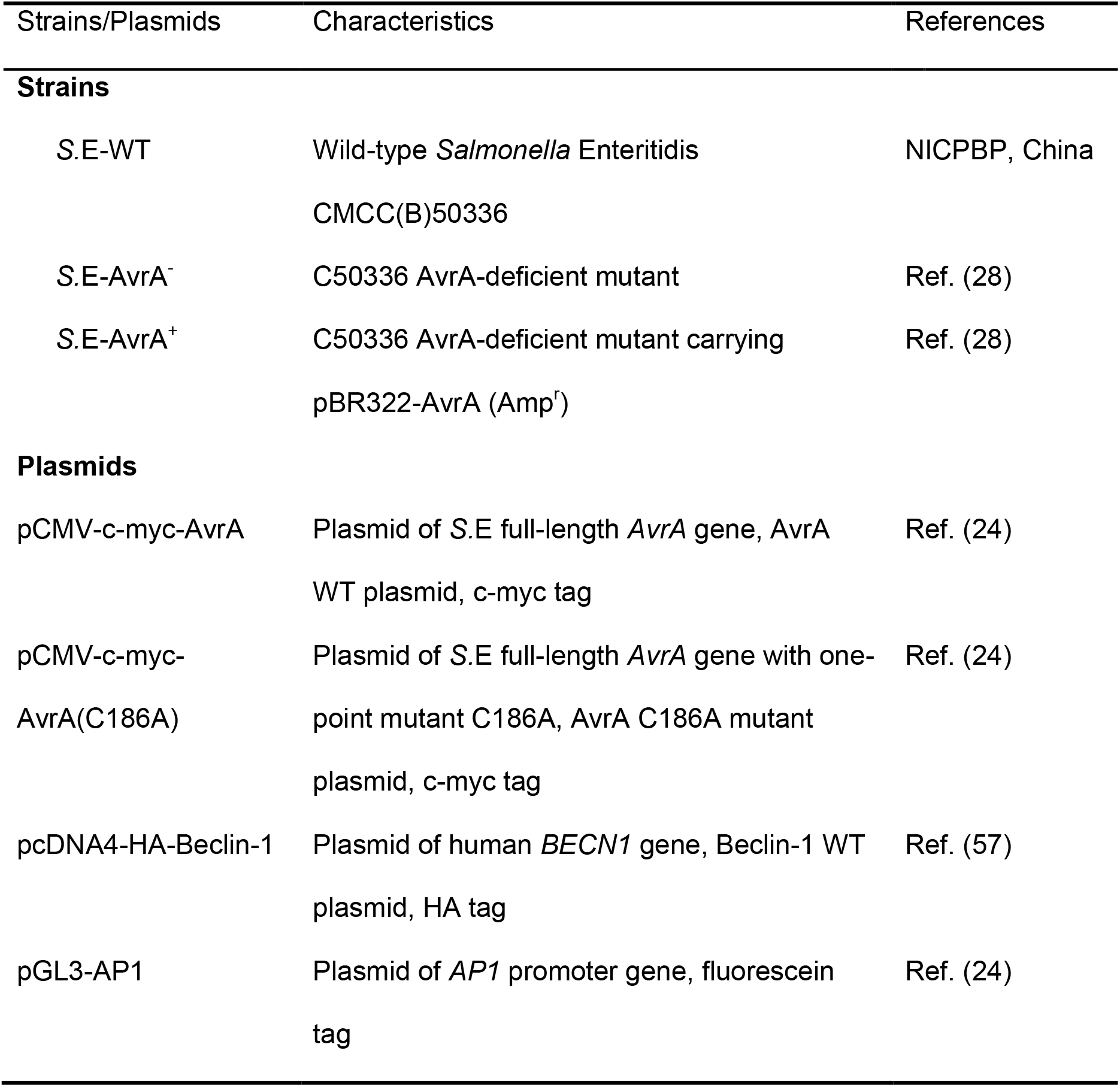
Bacterial strains and plasmids used in this study

### Cell culture

Human epithelial HCT116, Caco-2 BBE, and SKCO-15 cells were maintained in Dulbecco’s modified Eagle’s medium (DMEM) supplemented with 10% fetal bovine serum (FBS), streptomycin-penicillin and L-glutamine. Bacterial colony forming units (CFU) were determined by plating diluted cell lysates onto LB agar culture plates and incubating the cultures at 37°C overnight (28).

### Mouse intestinal organoid isolation, culture and passage

The mouse small intestines were removed immediately after cervical dislocation. The stool was then flushed out with ice-cold PBS (penicillin, 100 I.U./mL/streptomycin, 100 μg/mL), and the small intestines were dissected and opened longitudinally and cut into small (~1 cm) pieces. The tissues were rocked in PBS with 2 mmol/L ethylenediamine tetraacetic acid (EDTA) for 30 minutes at 4°C and were then switched to PBS with 54.9 mmol/L D-sorbitol and 43.4 mmol/L sucrose. The tissues were then vortexed for 1–2 minutes and were filtered through a 70 μm sterile cell strainer. The crypts were collected by centrifugation at 150 g for 10 minutes at 4°C. Approximately 500 crypts were suspended in 50 μL of growth factor-reduced phenol-free Matrigel (BD Biosciences, San Jose, CA). Next, a 50 μL droplet of the Matrigel/crypt mix was placed in the center well of a 12-well plate. After 30 minutes of polymerization, 650 μL of the mini gut medium was overlain (31, 51). The mini gut medium (advanced DMEM/F12 supplemented with HEPES, L-glutamine, N2 and B27) was added to the culture, along with R-Spondin, Noggin, and EGF. The medium was changed every 2–3 days. For passage, the organoids were removed from the Matrigel and broken up mechanically by passing them through a syringe and needle (27 G, BD Biosciences, San Jose, CA), and then, they were transferred to fresh Matrigel. The passage was performed every 7–10 days with a 1:4 split ratio.

### Bacterial colonization

Polarized human epithelial cells were grown in DMEM with 10% FBS. At 90-100% confluence, the cells were colonized with an equal number of the indicated *Salmonella* Enteritidis strain for 30 minutes, washed with Hank’s balanced salt solution (HBSS), and incubated in DMEM containing gentamicin (100 μg/ml) for 30 minutes. The first 30 minutes of the incubation allowed the bacteria to contact the epithelial cell surface and inject the effectors into the host cells (52, 53). After extensive HBSS washing, the extracellular bacteria were washed away. The incubation with gentamicin inhibited the bacterial growth (54). At the indicated times, post-colonization, the cells samples were harvested for the analysis.

The organoids (6 days after passage) were colonized with the indicated *Salmonella* Enteritidis strain for 30 minutes, washed with HBSS, and incubated in mini gut media containing gentamicin (500 mg/mL) for the indicated times, as described in our previous studies (31). After extensive HBSS washing, the extracellular bacteria were washed away. The incubation with gentamicin inhibited the growth of the bacteria. Samples were collected for a Western blot analysis or immunofluorescence after the organoids were colonized with *Salmonella* for 30 minutes and were then incubated in medium with gentamicin for 1 hour.

### Cell treatment with the JNK inhibitor SP600125

The JNK inhibitor SP600125 (50 mM, EMD Biosciences, San Diego, CA, USA) was added directly to the culture medium one hour before *Salmonella* treatment. The HCT116 cells were pretreated with SP600125 and were incubated with the indicated *Salmonella* for 30 minutes, washed 3 times in HBSS, incubated in DMEM containing 100 μg/ml gentamicin and 50 μM SP600125 for 30 minutes, and harvested. The protein levels were determined by Western blotting.

### Lysotracker staining

Lysotracker staining was performed following the manufacturer’s protocol (Life technologies). The HCT116 cells were grown in the Lab-Tek Chambered Coverglass System (154526, Thermo Scientific, Rockford, IL, USA), and the cells were then incubated with 100 nM LysoTracker Deep Red Probe (L12492, Life technologies, Eugene, OR, USA) in cell growth medium at 37°C for 60 minutes. After washing with HBBS, the cells were incubated with the indicated *Salmonella* for 30 minutes, washed 3 times in HBSS, incubated in DMEM containing 100 μg/ml gentamicin for 30 minutes, and the cells were detected by fluorescence microscopy (ECLIPSE E600, Nikon Instruments, Inc., Melville, NY, USA.).

### Streptomycin pre-treated Salmonella mouse model

Water and food were withdrawn 4 hours before an oral gavage with 7.5 mg/mouse streptomycin. Afterward, the animals were supplied with water and food. Twenty hours after the streptomycin treatment, water and food were once again withdrawn for 4 hours before the mice were infected with 1.0×10^8^ CFU of *Salmonella* (100 μl bacterial suspension in HBSS) or treated with sterile HBSS (control) by oral gavage, as previously described (32). At the indicated times post-infection, the mice were sacrificed, and the intestinal tissue samples were removed for the analysis.

### Immunoblotting and antibodies

Mouse epithelial cells were scraped and lysed in lysis buffer (1% Triton X-100, 150 mM NaCl, 10 mM Tris pH 7.4, 1 mM EDTA, 1 mM EGTA pH 8.0, 0.2 mM sodium orthovanadate, and protease inhibitor cocktail), and the protein concentration was measured using Protein Assay Kits (Bio-Rad, Hercules, CA, USA) (52). The cultured cells were rinsed twice in ice-cold HBSS and were lysed in protein loading buffer (50 mM Tris pH 6.8, 100 mM dithiothreitol, 2% SDS, 0.1% bromophenol blue, and 10% glycerol), and the remaining cells were scraped off the dish and sonicated to shear the DNA and reduce the sample viscosity. The organoid cells were rinsed three times in ice-cold HBSS and were then suspended in ice-cold HBSS. The organoid cells were then spun down at 900 rpm for 10 minutes at 4°C. Next, using a pipette to aspirate the PBS at the top, the organoid cells were lysed in lysis buffer and were then sonicated (31). The protein concentration was then measured. Equal amounts of protein were separated by SDS-polyacrylamide gel electrophoresis and were transferred to nitrocellulose membranes. The nonspecific sites were blocked with 5% bovine serum albumin (BSA) in TBST (50 mM Tris, 150 mM NaCl, and 0.05% Tween 20 adjusted to pH 7.6 with HCl), and the membranes were then incubated with dilutions of the primary antibodies as recommended by the manufacturers. The primary antibodies included the following: anti-p62 (1:1,000, AP2183B, ABGENT, San Diego, CA,USA); anti-LC3B (1:1,000, 2775), anti-SAPK/JNK (1:1,000, 9258), anti-phospho-SAPK/JNK (Thr183/Tyr185, 1:1,000, 9251), anti-c-jun (60A8, 1:1,000, 9165), anti-phospho-c-jun (S63, 1:1,000, 9261) (Cell Signaling, Beverly, MA, USA); anti-BECN1/Beclin-1 (1:1,000, SC-10086), anti-c-myc (PE10, 1:1,000, SC-40), anti-HA-probe (F-7, 1:1,000, SC-7392) (Santa Cruz Biotechnology, Inc., Santa Cruz, CA, USA); anti-β-actin (1:2,000, A5441, Sigma-Aldrich, Milwaukee, WI, USA); anti-ubiquitin (1:1,000, UG9511, ENZO, Farmingdale, NY, USA); and anti-AvrA (1:1,000, custom-made, as reported in our previous studies (55)). The membranes were washed and incubated with an HRP-conjugated secondary antibody (anti-mouse, 1:5,000, 31430; anti-rabbit, 1:5,000, 31460; anti-goat, 1:5,000, R-21459, Invitrogen, Grand Island, NY, USA). The membranes were then washed again, treated with the ECL Western Blotting Substrate (Thermo Scientific, Rockford, IL, USA) and visualized on X-ray film. The membranes that were sequentially probed with more than one antibody were stripped in stripping buffer (Thermo Scientific, Rockford, IL, USA) before re-probing.

### Immunoprecipitation

The cells were rinsed twice in ice-cold HBSS and were lysed in ice-cold immunoprecipitation buffer (1% Triton X-100, 150 mmol/L NaCl, 10 mmol/L Tris, pH 7.4, 1 mmol/L EDTA, 1 mmol/L ethylene glycol bis (β-aminoethyl ether)-*N,N,N’,N’*-tetraacetic acid, pH 8.0, 0.2 mmol/L sodium orthovanadate, and protease inhibitor cocktail (Roche Diagnostics, Basel, Switzerland)). The samples were prepared as previously described(56). The blots were probed with anti-HA-probe (F-7, 1:1,000, SC-7392, Santa Cruz Biotechnology, Inc., Santa Cruz, CA, USA) and anti-c-myc (PE10, SC-40, Santa Cruz Biotechnology, Inc., Santa Cruz, CA, USA) antibodies.

### Immunofluorescence

The ileal tissues from the distal portion of the ileum were freshly isolated and paraffin-embedded after fixation with 10% neutral buffered formalin. Immunofluorescence was performed on the paraffin-embedded sections (5 μm). After preparation of the slides, as described previously (53), the tissue samples were incubated with the indicated primary antibody, anti-lysozyme (1:100, sc27958, Santa Cruz Biotechnology, Inc., Santa Cruz, CA, USA), at 4°C overnight. The samples were then incubated with the sheep anti-goat Alexa Fluor 594 (A11058, Life Technologies, Grand Island, NY, USA) and DAPI (D1306, Life Technologies, Grand Island, NY, USA) for 1 hour at room temperature. The tissues were mounted with SlowFade (s2828, Life technologies, Grand Island, NY, USA), followed by a coverslip, and the edges were sealed to prevent drying. The specimens were examined with a Zeiss laser scanning microscope 710 (Carl Zeiss, Inc., Oberkochen, Germany).

### Paneth cell staining and counting

The Paneth cells in the mouse ileal tissue were counted after the anti-lysozyme immunofluorescence staining, according to our previous study(35). The patterns of the lysozyme expression in the Paneth cells were classified into four categories as follows: normal (D0); disordered (D1); depleted (D2) and diffuse (D3), according to previously published methods (36).

### AP-1 Transcriptional Activity Assay

The cells were transiently transfected with 1 μg of pGL3-AP1 plasmid using the Lipofectamine 3000 transfection kit, according to the manufacturer’s instructions (Invitrogen, Carlsbad, CA). The pRL-TK vector was used as an internal control reporter. After 24 hours post transfection, the cells were colonized with equal numbers of bacteria for 30 minutes, washed, and incubated in DMEM containing gentamicin (100 μg/ml) for 30 minutes. The luciferase activity was monitored using the dual luciferase assay system (Promega).

### Statistical analysis

Data are expressed as the mean ± SE. All statistical tests were 2-sided. The p values < 0.05 were considered statistically significant. The differences between two samples were analyzed using Student’s *t*-test; the differences among three or more groups were analyzed using one-way ANOVA. The Tukey’s method was used to adjust multiple comparisons to ensure results accurately. All statistical analyses were performed by GraphPad Prism 5(GraphPad Software, La Jolla, CA) or SAS version 9.4 (SAS Institute, Inc., Cary, NC, USA).

## Author contribution

Y. J., Z. L., Y. Z., C. M. and R. L.: data acquisition, analysis and interpretation, and drafting of the manuscript. Y. J., Y.Z., and J. S.: wrote the main manuscript text and prepared the figures. Y. X.: statistical analysis and drafting of the manuscript. Z. P., X. J. and X. X.: technical or material support and drafting of the manuscript. J. S.: study concept and design, critical revision of the manuscript for important intellectual content, and study supervision. All the authors reviewed the manuscript.

## Abbreviations

AvrA C186A mutation: mutated at the key cysteine required for AvrA activity
BSA: bovine serum albumin
CFU: colony forming units
DMEM: Dulbecco’s modified Eagle’s medium
EDTA: ethylenediamine tetraacetic acid
FBS: fetal bovine serum
HBSS: Hank’s balanced salt solution
IP: immunoprecipitated
JNK: c-Jun N-terminal kinase
LB: Luria-Bertani
*Salmonella* Enteritidis, *S*.Enteritidis, *S*.E: *Salmonella enterica serovar Enteritidis*
*Salmonella* Typhimurium: *Salmonella enterica serovar* Typhimurium
*S*.E-WT: *Salmonella* Enteritidis wild-type strain C50336
*S*.E-AvrA^-^: *Salmonella* Enteritidis AvrA deletion mutant
*S*.E-AvrA^+^: plasmid mediated complementary strain *S*.E-AvrA^-^/pAvrA^+^
SPI-1: *Salmonella* pathogenicity island 1
T3SS: type III secretion system
ub-Beclin-1: ubiquitinated Beclin-1

